# Genome-wide transcription factor activities are explained by intrinsic conformational dynamics of binding-sites and distal flanking-regions

**DOI:** 10.1101/020602

**Authors:** Munazah Andrabi, Andrew Paul Hutchins, Diego Miranda-Saavedra, Hidetoshi Kono, Ruth Nussinov, Kenji Mizuguchi, Shandar Ahmad

**Author notes:** Current address: RIKEN Center for Life Science Technologies (CLST), 2-2-3 Minatojima Minamimachi, Chuo-Ku, Kobe 6500047, Japan. Email addresses: Munazah Andrabi Andrew Paul Hutchins Diego Miraanda-Saavedra Hidetosho Kono Ruth Nussinov Kenji Mizuguchi Shandar Ahmad.

## Abstract

Transcription factors (TFs) recognize small DNA sequence motifs directly or through their sequence-dependent structure. While sequence composition and degeneracy are verified to be the defining factors of TF binding specificity, the role of conformational dynamics of the DNA remains poorly understood. With growing evidence from next generation sequencing (NGS) data suggesting the inadequacy of sequence-only models, alternative models for describing the TF binding preferences are required, wherein the conformational dynamics presents an attractive option. Here, we report a novel method (*DynaSeq*) which accurately predicts DNA-conformational ensembles for genomic targets of TFs. Using *DynaSeq* we demonstrate how the dynamics of binding sites and their distal flanking regions can be used to elucidate TF-binding patterns for two model systems: cell type-specific binding of STAT3 and chromatin structural specificity of 3 functional TF classes viz. pioneers, settlers and migrants. We find that TF preferences in both these systems can be accurately explained by the conformational dynamics of their binding sites and their distal flanking DNA regions. Conformational dynamics not only distinguishes binding sites from genomic backgrounds in STAT3; it also points to a modular organization of their surrounding regions. Further, the differential binding modes of STAT3-DNA reveal a potential mechanism of cellular specificity. Our model identifies clear signatures to accurately classify pioneer, migrant and settler TF targets from the dynamics of distal flanking regions. This suggests that the chromatin preferences of TFs are significantly influenced by the intrinsic conformational dynamics of the DNA surrounding the TF binding sites.

## Introduction

The physical basis of protein-DNA interactions has thus far been investigated from the perspective of *direct recognition* of nucleic acid bases by complementary TF residues or through an *indirect recognition* of sequence dependent DNA structure[1-4]. While the former ignores the differential accessibilities of DNA bases in the double helix the latter assumes the existence of a unique and exclusive structure of the DNA. We have previously, developed techniques to thread DNA sequences onto the structure of a known protein-DNA complex and determine the energy of native and designed sets of sequences[5-11]. Using these predicted intrinsic DNA energies, sequence specificity of the DNA structure in a given protein-DNA complex was successfully explained. However, focused on the assumption of an energetically most favorable structure, none of these methods addresses the *dynamics* of the molecules. Since the system has been assumed to be rigid the role of conformational dynamics in shape component or indirect readout mechanism of protein-DNA recognition has been overlooked.

In addition to the assumption of a static structure of the binding site the role of its flanking regions and their structure at a genomic scale also remains unexplored. Since most high resolution studies on this subject rely on crystal structures of TFs bound with small DNA fragments (typically just 7-8 bp DNA), structural information away from the binding sites are unavailable and commonly ignored[12]. Yet, structural patterns around the binding sites may be crucial for cellular and *in vivo* (versus *in vitro*) TF binding specificities [13]. It is therefore important to develop a method to investigate the sequence-dependent DNA-conformational dynamics at a genomic scale and examine its role in binding site recognition.. Here, we present a novel approach, called *DynaSeq* to model DNA sequence specificity by sequence-dependent conformational ensembles. *DynaSeq* is a set of support vector regression models (SVMs), trained over molecular dynamic trajectories of tetranucleotides and thoroughly benchmarked by cross-validation during the training of the model followed by comparison of the predictions with structures in protein data bank (PDB).

We investigated the role of conformational dynamics in TF binding by applying *DynaSeq* to two model systems. In the first case, the DNA conformational dynamics of genome-wide binding sites of STAT3 reported in four distinct cell types[14] were predicted and analyzed. We report that the DNA conformational dynamics successfully explains observed genome-wide STAT3 binding data with high accuracy, without explicit use of sequence information. Further, *DynaSeq* reveals distinct blocks of information-rich regions whose importance to TF binding falls off at different rates. Conformational dynamics profiles of sequence data as well as the consensus DNA conformation alignments with PDB structure reveal cellular-specificity and a potential mechanism of recognition.

We also looked into the patterns of TF recognition in the chromatins using *DynaSeq*. Recently TFs have been characterized as *pioneer, migrant and settler* factors *[15]*. We investigated if these three groups of TFs recognize similar DNA structures and whether the members of each group can be predicted directly from the DNA conformational dynamics of their binding sites and/or flanking regions.

Analysis of putative binding sites within 200 base pairs stretches in both the above systems indicates that sequence regions flanking the core binding sites are highly informative for TF binding preferences. These findings are significant because they may provide a missing link between condition-specific TF binding and the description of cellular environments. Taken together, this study provides a novel approach to study DNA structural dynamics at a genomic scale and indicates that information about TF-DNA binding is contained not only in the exact site of TF-binding but also extends to much larger flanking region of DNA.

## Results

*DynaSeq* takes a DNA sequence as input and for each base position it predicts a 60-dimensional ensemble (12 conformational parameters reported by the 3DNA program[16] with 5 bins each) (See Methods). *DynaSeq* was benchmarked and employed to investigate the conformational ensembles of STAT3’s genome-wide binding sites and to distinguish between *pioneer, migrant* and *settler* TFs.

### **Development and benchmarking of *DynaSeq*:**

*DynaSeq* was trained on Molecular dynamic (MD) simulations data of 136 unique tetranucleotides, represented by 60-dimensional conformational ensemble for each nucleotide position (see Supplementary Methods SM1-SM7; Supplementary Results SR1; Supplementary Table ST1). To evaluate *DynaSeq*, we created independent training and test sets in a leave-one-tetranucleotide-data-out fashion; train the ensemble populations for all base positions in 135 tetranucleotides and test the predictive power for the left-out 136^th^. Results from an exhaustive set of 136 combinations were pooled and evaluated.

The predictive model in each case is a set of 60 SVMs (one SVM for each ensemble dimension), where the inputs to the SVM are identities (A, C, G or T) of a DNA base and its sequence neighbors within a window and the outputs are the corresponding ensemble populations. Figure 1 summarizes the prediction performance of these cross-validated models. We trained and tested various window spans and found that a 5-nucleotide window is optimum for the prediction model. Based on this optimized model, most of the populations are well predicted (∼87%) with an absolute error of 5 percentage points (global mean for each bin is 20%) (Figure 1b). A high correlation (R=0.88) between the predicted and observed values in the entire population range further indicates the stability of the prediction model (Figure 1c). Furthermore Figures 1(c) and (d) indicate that the populations, highly skewed from their global 20% value are estimated well albeit with a slightly higher error rate than for the values in the middle. However, the worst-case MAE is only about 6% (observed in the first bin of the *helical twist*) providing confidence for its genome-wide applicability.

**Figure 1.**
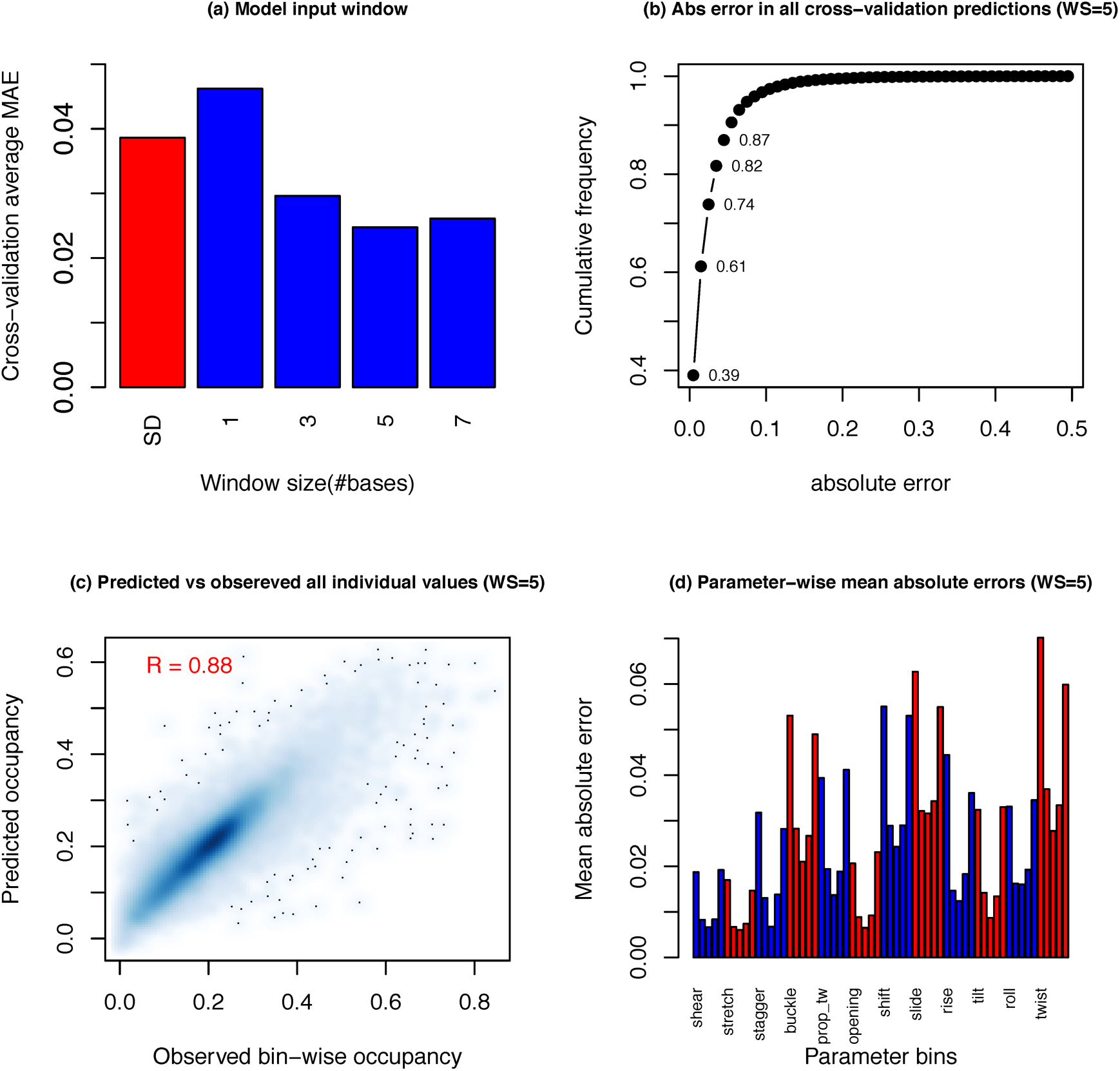
Cross-validation predictability of DNA conformational ensembles at each base position (a) variation of mean absolute error (absolute difference between prediction and observed population density in each bin) with training window size. Standard deviation in the overall data is shown in red, whereas other values represent cross-validation performances (b) Overall cumulative frequency of absolute error distribution at window size=5. Prediction for each base in any position of a tetranucleotide is counted once and errors computed are for the left out sets of leave-one-tetranucleotide cross-validations. First few populations are labelled. (c) Scatterplot of predicted versus observed population in all bins and all conformational parameters (d) Mean absolute error averages within each bin of all parameters.

We also evaluated *DynaSeq* as a predictor of static DNA structure in addition to dynamics utilizing solved crystal structures in Protein Data Bank (PDB). The results indicate that real DNA structures could be favorably modeled from their predicted dynamics with an RMSD of 3.74Å compared to 6.28Å for randomly generated sequences. This result also corresponds to a negative Z-score of 1.38 (all median values used). (Supplementary Results SR2, and Supplementary Tables ST2 and ST3). These results provide additional support for *DynaSeq* applicability on a blind data set.

Thus, *DynaSeq*, which can capture sequence-dependent patterns of conformational dynamics, is a powerful tool providing useful insights into TF DNA binding. Furthermore, owing to its low computational cost *DynaSeq* makes its highly feasible to investigate the conformational dynamics of the high throughput genomic data being produced at an unprecedented scale.

Our aim is to obtain biological insights into the genome-wide TF recognition process using conformational dynamics through *DynaSeq*. Even though some of the information that *DynaSeq* produces may also be obtained from *DNAShape [2]*, there are significant differences, as the latter cannot produce all the information provided by *DynaSeq*. Firstly, *DNAShape* gives only four structure-based parameters and hence does not provide comprehensive prediction of DNA structure. More importantly, conformational ensembles can exclusively be computed by *DynaSeq* hence making a comparison between structure and dynamics possible. On the technical side, conformations used in *DNAShape* are derived from *Monte Carlo* simulations whereas *DynaSeq* is based on MD, even though it is not clearly know which proves to be a better predictor for DNA structure. Nevertheless, we compared the four predicted parameters of *DNAShape* with all the 60 ensemble bin population predictions of *DynaSeq* (and also the 12 parameters denoting its averaged structure) for all unique DNA 6 mers. Even though the conformational parameter definitions do not have an exact correspondence between the two sets, we observed good agreement between several pairs particularly the propeller twist from both methods and other pairs involving the DNAShape’s minor groove width (Pearson correlation exceeding 0.4 in both cases) suggesting a reasonable degree of consistency between the two approaches (Detailed comparisons in Supplementary Table ST4). In summary, even though some similar information on DNA descriptors can be obtained by *DNAShape*, *DynaSeq* makes possible a more thorough analysis of both structure and dynamics.

### DNA conformational dynamics in STAT3 target sequences

STAT3 is an important TF with numerous cellular functions. [17]. It is a key regulator of immune response, interacting with other key factors leading to its phosphorylation, dimerization and translocation to the nucleus, where it binds the promoters of downstream host defense genes, regulating their expression. The mechanism by which STAT3 recognizes its DNA targets is not well described and hence its binding to DNA under different cellular contexts is nearly indistinguishable [14, 18]. Even though a canonical sequence motif (TCCnnnGAA) has been reported, it appears to be non-essential for STAT3 binding[14]. The absence of a core sequence motif in all STAT3-bound genomic targets prompts us to look for an alternative model of TF recognition and DNA conformation provides an excellent opportunity to do so. Interestingly, the role of DNA shape in distinguishing STAT1 and STAT3 binding mode to its target has been recently suggested[19] further strengthening our idea of a using conformational dynamics for target selection. We recently discovered a novel non-canonical target of STAT3 in *Saa* gene with the help of our DNA-structure threading method *Readout* [7, 20]. To investigate the DNA-conformational dynamics of STAT3 targets, we exploited its genome-wide binding site data in four distinct cell types available in the public domain [14, 18, 21]. Four cell types namely (1) Embryonic Stem Cell [22] (2) CD4+ T cells [23] (3) AtT-20 corticotroph cells [24] and (4) macrophages [25] for which ChIP-Seq data are available were analyzed. These ChIP-Seq experiments for all four cell-types have been performed independently, addressing specific questions focused in corresponding publications. We have earlier reported an integrated analysis of these results [14] and compiled a set of STAT3 from the cell-type specific data. The same targets have been reused in this work for a genome-wide conformational dynamics analysis as per the protocol summarized in Figure 2. General and cell-specific conformational dynamics patterns, revealed from this analysis are presented in the following.

**Figure 2.**
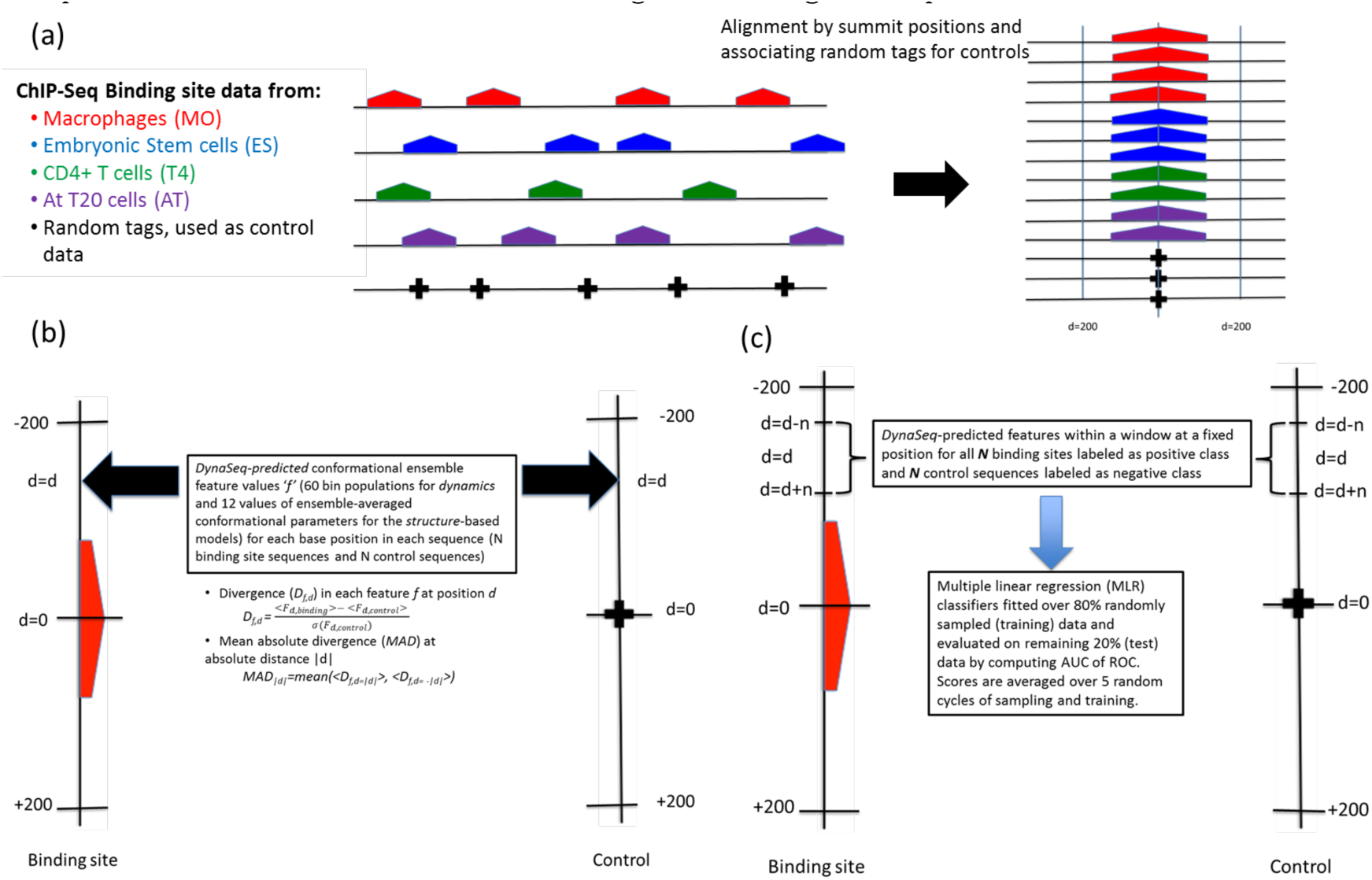
Flowchart for the analysis of genome-wide STAT3 binding sites. (a) ChIP-Seq peaks are first aligned by their summit positions and a data set of genomic sequences around summit positions (200 bases on either side) is compiled. Equal number of randomly tagged genomic sequences is also collected to serve as control. (b) *DynaSeq* is used to predict conformational ensemble for each position in each binding site and control sequences. This returns a 60 dimensional feature vector representing conformational ensemble (dynamics model) at each position in each sequence. Averaged values of 12 conformational parameters (structure model) are also obtained. Feature values between control and binding data are compared at each position individually (feature divergence) and as an average of all features (Mean absolute divergence or MAD). (c) Multiple linear regression models are created to assess the ability of computed features within a window to distinguish binding site sequences from controls.

#### Overall conformational dynamics of genome-wide binding sites

As illustrated in Figure 2, we aligned stretches of putative binding sites reported from ChIP-Seq data by their *summit* positions and analyzed the patterns of conformational dynamics in contrast to random genomic backgrounds at various alignment positions. We computed the Mean Absolute Divergence (MAD) of conformational ensembles, which quantifies how the individual alignment positions of binding data differ from the controls over a large distance range, in terms of conformational dynamics-derived features (Figure 3(a)). MAD scores indicate binding site regions surrounding the summit to be composed of four distinct regions such that the extent of conformational divergence within each region follows a specific pattern. The first, ∼ 19-nucleotide window *Region-I*, (summit +/-9 bases) has the expected highest MAD scores as this is the region, where the TF is presumed to directly bind its anticipated sequence-specific binding site. We observe that the divergences at the summit positions are smaller than the adjacent regions (positive slope in Region-I), along expected lines as the summit position of STAT3 binding region corresponds to the linker region of its canonical binding motif, whereas the actual binding regions are a few bases apart [14]. More interestingly, there appears a *Region-II* (10^th^ to 50^th^ base positions from the summit), in which the divergence variation is almost constant (the slope of the linear regression curve within this region is 0.0062, which is about 1/3^rd^ of the slope of the regression curve of the next region) indicating an equal contribution to TF recognition from all the bases within this region. In the *Region-III* (position 50-100), the information falls off almost linearly and suddenly drops at its boundary with the next region (d∼100). In the last region (*Region-IV*), the distribution becomes flat again (slope falling to about half as compared to the previous region), slowly converging to a background value (Figire 3(a)). A non-zero background value is indicative of random variations and provides a good reference point for other meaningful positions.

**Figure 3.**
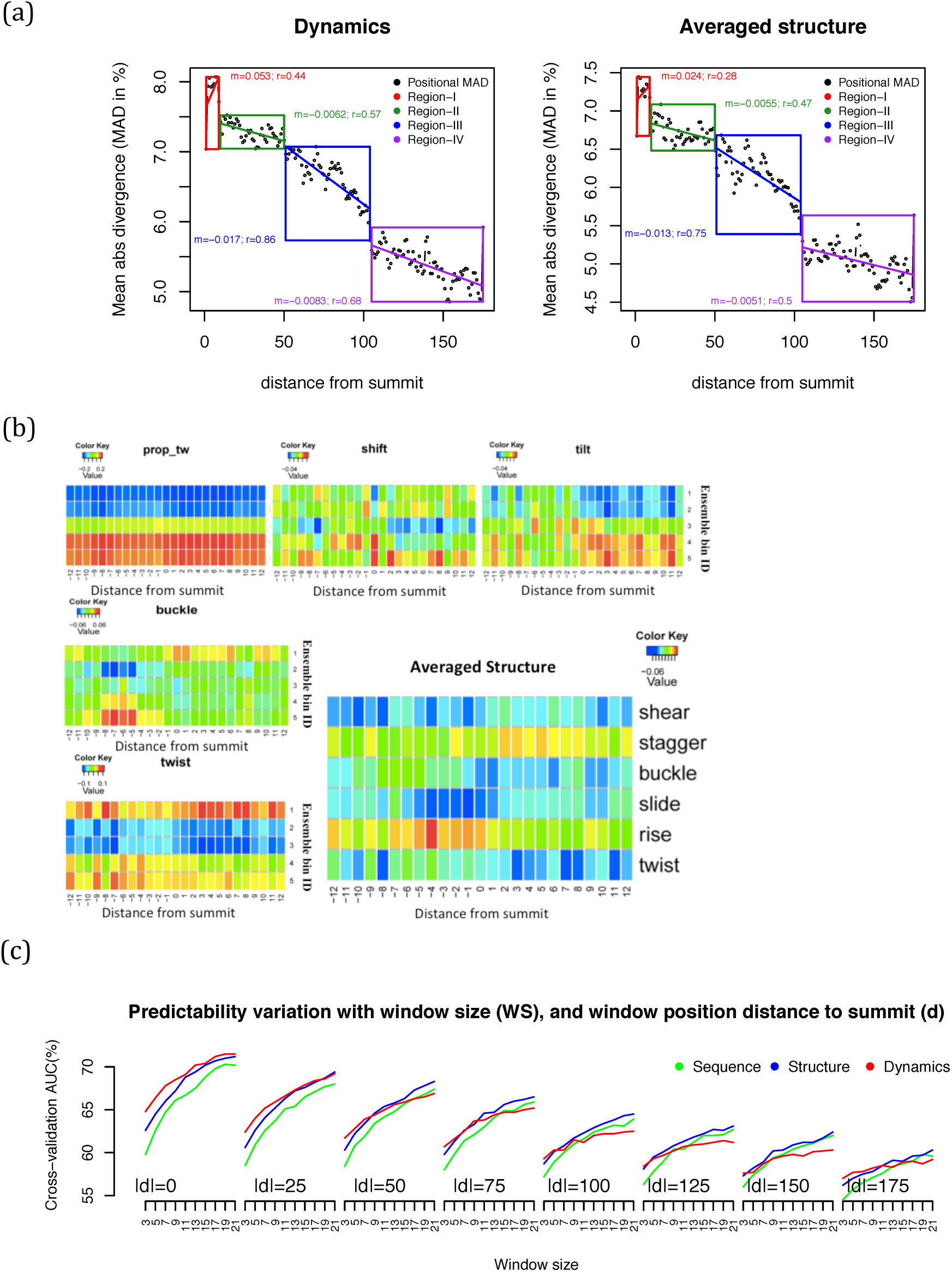
Enrichment of conformational ensemble features in STAT3-binding sites compared to genomic controls: (a) Variation of Mean absolute divergence (MAD) with distance from summit position (moving average of 3-base window), indicating how different is the DNA structural dynamics profile in binding site sequences compared to control data. Sequence regions as far as 200 bases from the summit positions show sufficient enrichment in each ensemble populations. MAD scores show sudden drops at about 18, 50 and 100 base positions from the summits, suggesting that the genomic sequences form distinct structural blocks each with a potentially different role in nucleotide positions in the sequence. Four contiguous regions are labeled and piece-wise linear regression plots are used to show that the slopes (m) within each region are distinctly different and that the dynamics fit regression model with better correlations (r) in each region.(b) Conformational variations at short distances from the summit positions. Some parameters show a constant position-independent variation indicating general compositional bias, whereas others form a clear structural signature (e.g. higher of tilt at specific 5’ positions and a clear pattern of high buckle in -6 to -9 base positions). Many of the conformational populations are also reflected in their predicted average structures. (c) Ability of cross-validated linear regression models to distinguish genome-wide STAT3 target sites from genomic backgrounds, and thereby explain the results of ChIP-Seq experiments. Similar regression models of various window sizes are created from sparse-encoded sequence, predicted conformational ensembles and predicted averaged structure. Regression model performance at position *“d”* and *“–d”* are computed in a cross-validation manner and the AUC of ROC indicating predictability of models to identify unseen binding sites from the two models are averaged. These plots show that the sequences far from summit position contain significant structural information, which may be crucial for transcription factor activity.

For this particular plot, similar patterns are observed in the dynamics and the averaged structure. However, Pearson correlations between predicted values from fitted linear models in each region are consistently higher, which suggests that the former is a more accurate reflection of TF-binding activity. Since all these correlation coefficients are reasonably high (0.44 to 0.86 for the dynamics model) their contributions to TF activity are statistically appealing.

We next analyzed the conformational dynamics in and around the neighborhood of *Region-I* by plotting divergence scores of individual conformational parametres for a 25-nucleotide window placed at the summit position. Here, we show plots from a selected number of conformational parameters *prop_tw, shift*, *tilt* and *buckle* and *twist* dynamics and for selected parameters for the averaged structures (Figure 3(b), complete set of heatmaps for the divergence scores for the 12 conformational parameters are included in Supplementary Figure SF4). We observe that certain features (predicted ensemble population in a bin for a given conformational parameter such as *prop_tw slide, opening, roll, stretch* and *rise)* show a largely constant divergence within the distance range considered. Higher value of *rise* is generally indicative of the elongation of minor or major *groove*, which results in a possible increase in chromatin accessibility, and its prominence in our data is consistent with the previous reports on their role in DNA recognition[26, 27]. However, this type of position-independent divergence of ensemble populations could simply be an sign of compositional biases of nucleosome-free or of other regions involved in DNA-binding and hence the divergences in certain positions for several other conformations such as *shift*, *tilt*, *buckle* and *twist* are probably more meaningful for specificity. For example, -5 to -8 base positions are characterized by high *buckle* conformation, whereas positions +2 to +12 are more informative in terms of *shift* and *tilt* parameters. The enriched bins near the summit positions for some of the parameters (*slide and twist*) are not skewed in just one direction but are separated by a disfavored bin between two or more enriched bins. This observation is unique to few parameters as the conformational features of most other parameters show a systematic shift only in one direction. The two distinct conformations for these parameters seem to form stable STAT3 target complex indicating the possibility of multiple and compound binding patterns. The complex statistics of these conformational parameters and their clear role in target recognition by TFs may possibly be utilized in designing high affinity targets and DNA inhibitors, not investigated in this work.

To estimate the collective role of several neighboring bases for distinguishing binding site sequences from controls, we developed simple multiple linear regression (MLR) classifiers using *DynaSeq*-predicted ensemble populations as the model inputs. The prediction performance of these models is a good measure of binding site information contained in a certain window (size=W). To estimate whether the ensemble populations (60×W-dimensions) are more (or less) informative than sequence information (4×W-dimensions) or the *DynaSeq*-derived averaged structure (12×W-dimensions), similar models were created for them as well. Several *cross-validation* models were developed by training 80% of the sampled data, testing the remaining 20% and averaging the performances from these cycles. We point out that cross-validation is somewhat ‘unfair’ to the dynamics-based models compared to sequence and structure (due to a larger number of features that could lead to over-fitting on training data, thereby reducing the predictive ability on the test samples), especially for the larger window sizes. Nonetheless, if we can show that the dynamics models perform better or equally well compared to sequence and structure based models, their superiority would be convincingly established. Figure 3(c) shows the results obtained from these cross-validated models. It becomes clear that at the summit and nearby positions, the dynamics based models perform better than sequence and structure; especially at small window sizes (small windows are less likely to be over-trained and reflect a more realistic scenario for comparison). At the very least, we have clearly established that the structure and dynamics carry all the information contained in the sequence (in a latter section, we convincing establish that dynamics is consistently more informative than a static structure in its predictive power). Interestingly, the sequence-encoded DNA conformational dynamics remains informative more than 100 bases away from the summit position, which is also revealed by the MAD score presented in Figure 3(a). Finally, more than 70% of the binding site data could be distinguished from control sequences using dynamics alone at the summit positions, which is clearly better than sequence-based PWMs as reported earlier[14].

#### Conformational ensemble and cellular specificity

In order to determine the relative effectiveness of sequence, structural and dynamics-based features in modeling the cellular specificity of STAT3 targets, we repeated the above MAD analysis for cell-type specific binding sites. Statistical details of these analyses are shown in Supplementary Figure SF5. We observed that the boundaries of the four regions identified above are more or less retained in at least two of the four cell-type specific data sets, macrophages and CD4+ T cells. However, the pattern of information distribution is vastly different in each. For example, the Region-II and not the Region-I is surprisingly more informative in macrophages unlike other cell-types. In the case of CD4+ T cells, Region-III is more informative than Region-II and the information drop at the start of Region-IV is the sharpest of all cell types. In the case of ES and AtT-20 cell types, the information falls off from Region-I to IV almost linearly, suggesting that the four regions almost merge with each other. The exact reason of such variations in the information storage patterns is not clear, but it is remarkable that the DNA conformational dynamics carry a cell-specific signature for up to a couple of hundred bases from the summit, potentially playing a critical role in cell type specific patterns of gene expression.

#### STAT3 binding centered on STAT3 DNA motif location

The results above provide useful clues into the general and tissue-specific recognition of genomic targets by STAT3. To gain further confidence in these results, we attempt to rule out the following potentially confounding factors: (a) Typical ChIP-Seq data have low base pair resolution and the precise location of STAT3 binding sites remains ambiguous. Therefore, some actual binding sites are likely to be present away from the summit positions. Although this bias cannot be completely eliminated, we believe that a significant enrichment of binding sites beyond a certain distance from summit position is highly unlikely [25]. Yet, a dataset accurately aligned by their precise binding sites will be helpful in establishing beyond doubt, if the binding site distal regions contribute to STAT3 recognition, as proposed above (b) We compared in the previous sections, the ability of the sequence, structure and dynamics to explain the STAT3 binding sites by training MLR models and used cross-validation strategy to eliminate the undue advantage to higher dimensional models (e.g. dynamics versus sequence (Figure 3c). This in turn gives advantage to lower-dimensional models, due to an over-fitting on the training data for higher dimensional models. A thorough distinction between the effectiveness of sequence, structure and dynamics based models can be established if the three feature sets in them could be uniformly represented.

We reanalyzed our data presented in the previous sections by addressing these two issues simultaneously. We created a subset of our ChIP-Seq reads by identifying the motif positions and aligning them by these motif centers, rather than the peak summit. This process of subset selection is disadvantageous as not all STAT3 binding sites actually contain a STAT3 DNA binding motif [14], and such STAT3 bound DNA having low quality motifs will be discarded. Indeed only a small fraction of ChIP-Seq reads were found to contain the detected sequence motif as shown in Table 1. Nevertheless, it will provide an unambiguous understanding of the patterns of information contained in regions free from actual binding sites. To analyze this motif-centered data, we computed principal components (PC) of the structure and dynamics representations for all possible 5-mer sequences (see Methods), which allowed us to compare the three feature sets under uniform dimensionality. We selected three sets of principal components (PC1-4, PC5-8, PC9-12) to create MLR classifiers similar to previous sections, and investigated their performance at the precise binding sites and in their neighborhood by placing a fixed window at these positions. Figure 4 shows the results of both (a) and (b) and provides powerful insights into the way conformational ensembles can be exploited by TFs for target recognition.

**Figure 4.**
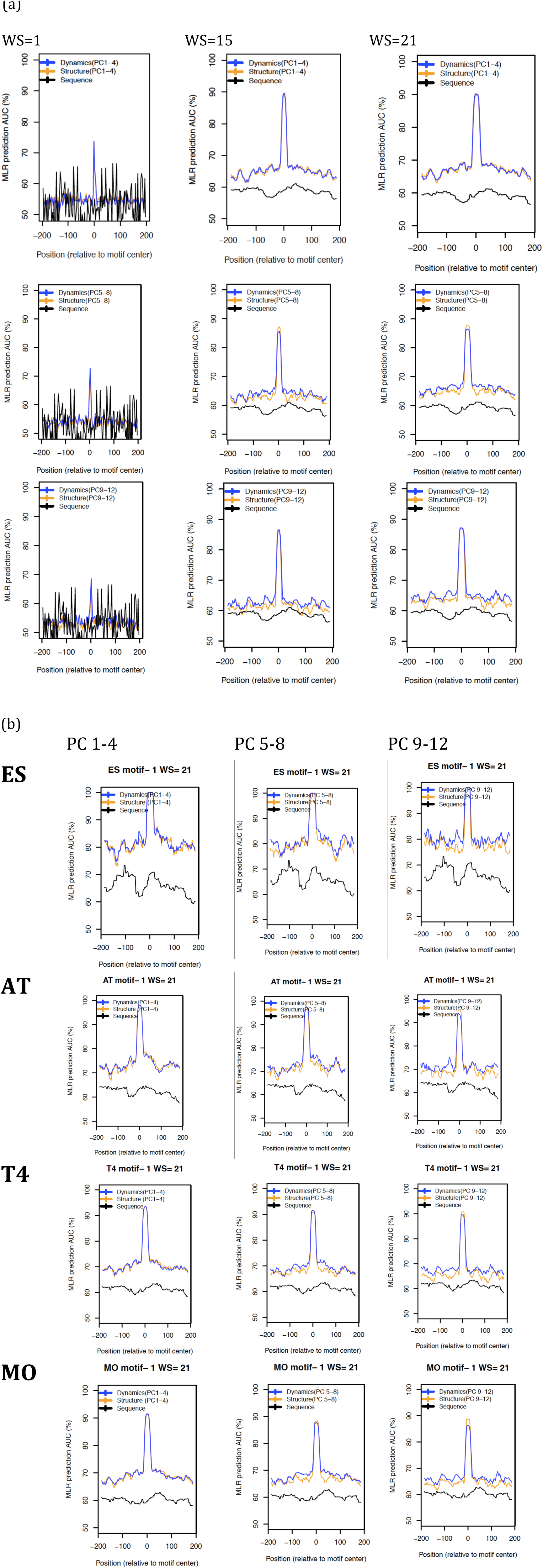
Specificity of STAT3 binding sites aligned by MEME-detected sequence motifs in terms of information contained in a set of four principle components. (a) Three motifs were detected by MEME in the overall data and sites containing these motifs are used to develop self-trained MLR models using differently defined 4 features derived from sparse-encoded sequence, sets of four principal components (PC1-4, PC5-8 and PC9-12) from structure and dynamics based ensembles Sequence motifs similar to (a) derived for STAT3-sites for each motifs and sequences collected for each cell-type independently. Structure is found to provide long-range cooperativity between nuclic acid sites (supported by the observation of higher performances on larger windows), and dynamics-based ensembles easily outperforms averaged structure-based models establishing the significance of conformational ensemble in addition to a static structure. Predictability of STAT3 sites at distances away from motif center support the conclusions derived from preceding sections (see manuscript text).

**Table 1.**
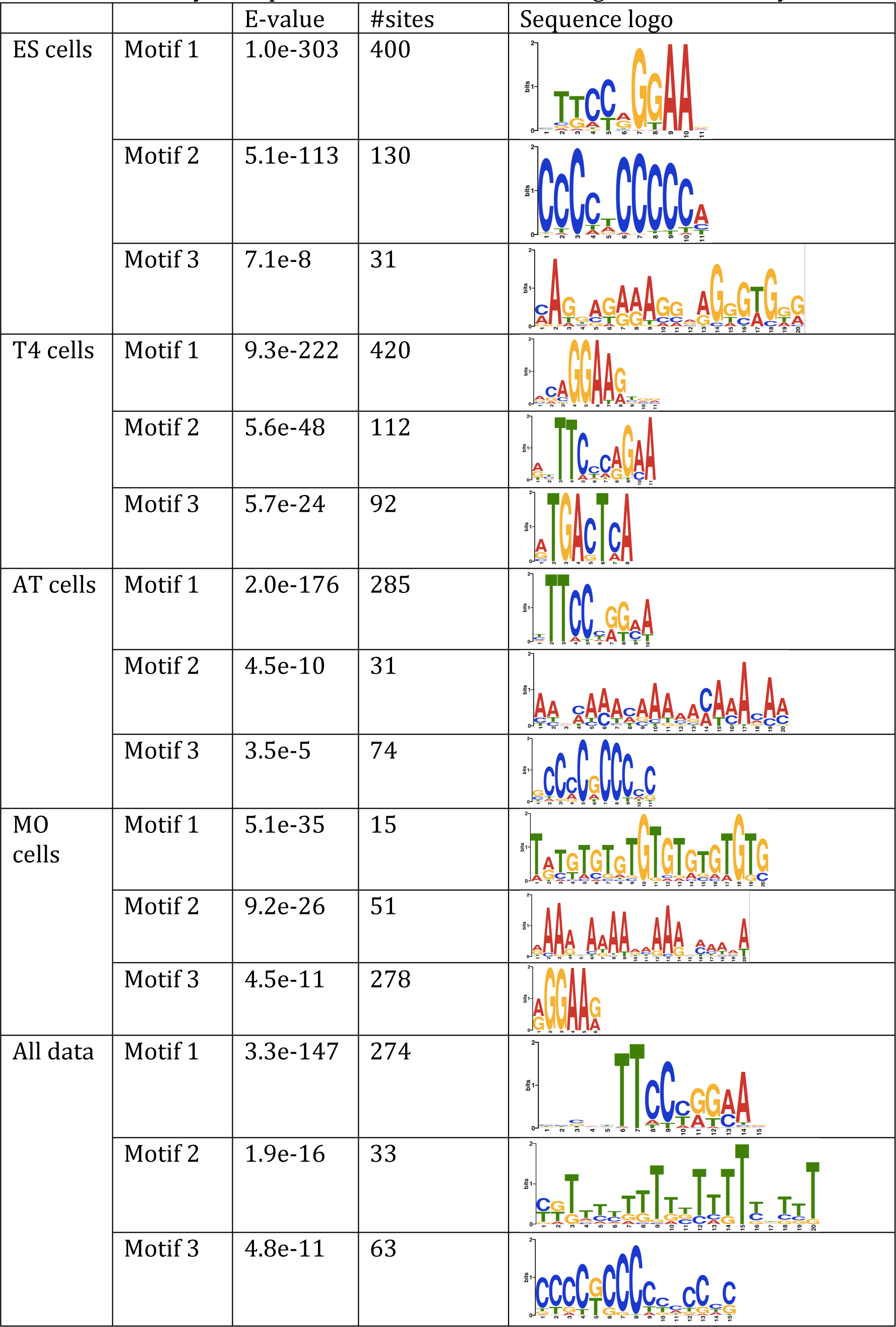
Summary of sequence motifs in STAT3 binding sites detected by MEME.

|  |  | E-value | #sites | Sequence logo |
| --- | --- | --- | --- | --- |
| ES cells | Motif 1 | 1.0e-303 | 400 |  |
|  | Motif 2 | 5.1e-113 | 130 |  |
|  | Motif 3 | 7.1e-8 | 31 |  |
| T4 cells | Motif 1 | 9.3e-222 | 420 |  |
|  | Motif 2 | 5.6e-48 | 112 |  |
|  | Motif 3 | 5.7e-24 | 92 |  |
| AT cells | Motif 1 | 2.0e-176 | 285 |  |
|  | Motif 2 | 4.5e-10 | 31 |  |
|  | Motif 3 | 3.5e-5 | 74 |  |
| MO cells | Motif 1 | 5.1e-35 | 15 |  |
|  | Motif 2 | 9.2e-26 | 51 |  |
|  | Motif 3 | 4.5e-11 | 278 |  |
| All data | Motif 1 | 3.3e-147 | 274 |  |
|  | Motif 2 | 1.9e-16 | 33 |  |
|  | Motif 3 | 4.8e-11 | 63 |  |

We observed that at single base resolution, sequence, structure and dynamics present a similar level of information as their capacity to distinguish between STAT3 binding sites from genomic sequences is comparable as seen from results for a single base window size (WS=1 in Figure 4(a)). However, the gap between structure and dynamics versus sequence widens indicating that the former have a higher predictive power than the latter. The structure captures the cooperativity between the TF and its target bases, whereas the sequence works largely at single base resolution. Secondly, the performance with the first four principle components of structure and dynamics are almost identical and hence dynamics provide no advantage over structure in these components. However, in higher components (PC5-8 and higher; second and third row of Figure 4(a)), dynamics-based MLR models clearly outperform structure-only models. Thus, the conformational dynamics predicted by *DynaSeq* is a better representation of TF binding sites than sequence or structure alone. Third the possible existence of three regions of ∼50-base each is broadly supported by plots in Figure 4(a) and (b). We do observe secondary and tertiary peaks in the MLR model performances around the same intervals as reported above. However, clear distinctions of the three regions, outlined by MAD in Figure 3 are blurred in some of these plots, which is likely due to multiple sources of noise in these positions and the fact that we are working on a smaller data set of only the sequences containing a sequence motif We also confirm the predictive power of regions far from the motif center, which is encoded in the sequence-dependent structure and dynamics. Finally, Figure 4(b) supports that the conformational ensemble profiles as being tissue-specific as the shape and locations of secondary peaks in the MLR model show subtle differences in all principal components examined here.

#### Comparison with the crystal structure of the STAT3/DNA complex in the PDB

Mouse STAT3 dimer structure in complex with DNA has been solved by Xray crystallography at 2.5Å resolution[28]. To gain insights into the cell-type-specific mechanism of complex formation, we compared the *DynaSeq*-derived averaged conformational parameters from each cell type with those in the crystal structure of the DNA in this complex. We created DNA sub-structures from the complex and aligned these “structure motifs” with consensus signatures derived from *DynaSeq*-predictions for the genomic data of each cell type (see methods). Curiously, ChIP-Seq summit positions from each cell type were found to align differently when a comparison between predicted structure for binding sites from a cell-type and the observed structure in the crystal structure was performed i.e. the best structural alignment was found to be slightly different sequence positions in different cell type data. Preferred “structural motifs” (here represented in terms of the DNA-sequence fragments in the PDB) from each cell type are shown in Figure 5a and the mapped alignments are shown in Figure 5c and Figure 5d. Macrophage targets of STAT3 are found to be unique in terms of size, location and the fact that only two (very similar) alignments were observed. The summit position from macrophage targets aligns best with the central base in the first TTT occurrence in the PDB structure (6^th^ base from the terminal), seemingly indicating a STAT3 half-site, something seen commonly in macrophages as compared to the other cell types[14]. On the other hand, target data from ES cells and At-T20 cells share their top two alignments, a trinucleotide (CCC) at the 9^th^ base position and CGT at the 11^th^ base position. Target data from CD4+T cells aligned best around position 8, overlapping with the well-known half-site of the canonical motif (TCC). Longer alignments around the same position are also observed for the same cell-type data. Thus, when mapped to their crystal structure, we observe that the conformational ensembles at the summit position resemble slightly different sites. The data does indicate that there is a small shift between the exact summit positions with respect to the DNA structural elements required for binding, pointing to cellular specificity of information-rich regions (Supplementary Figure SF5). The crystal structure data also suggest that there are higher resolution changes in protein-DNA interactions across cell-types, requiring further investigation.

**Figure 5.**
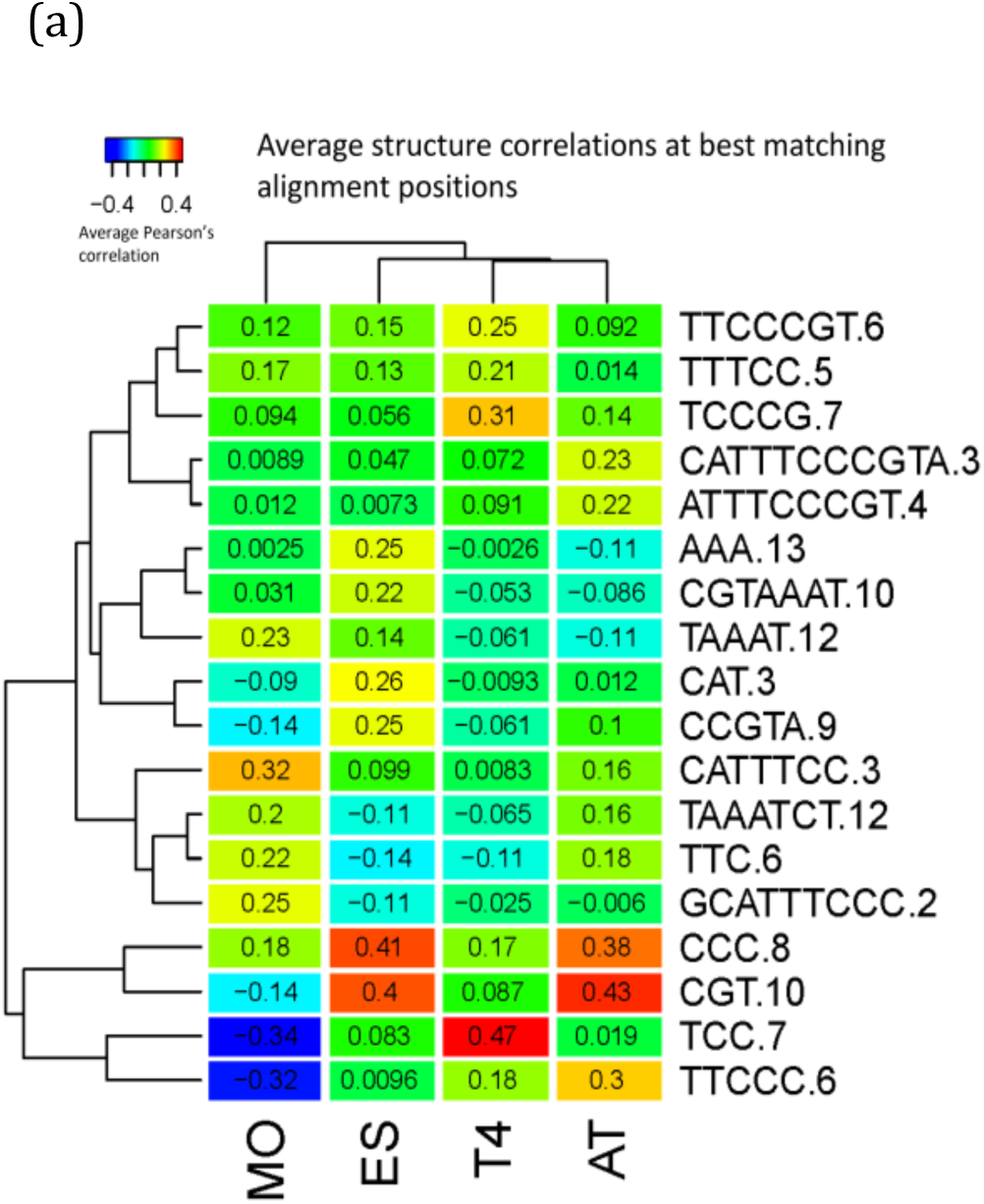
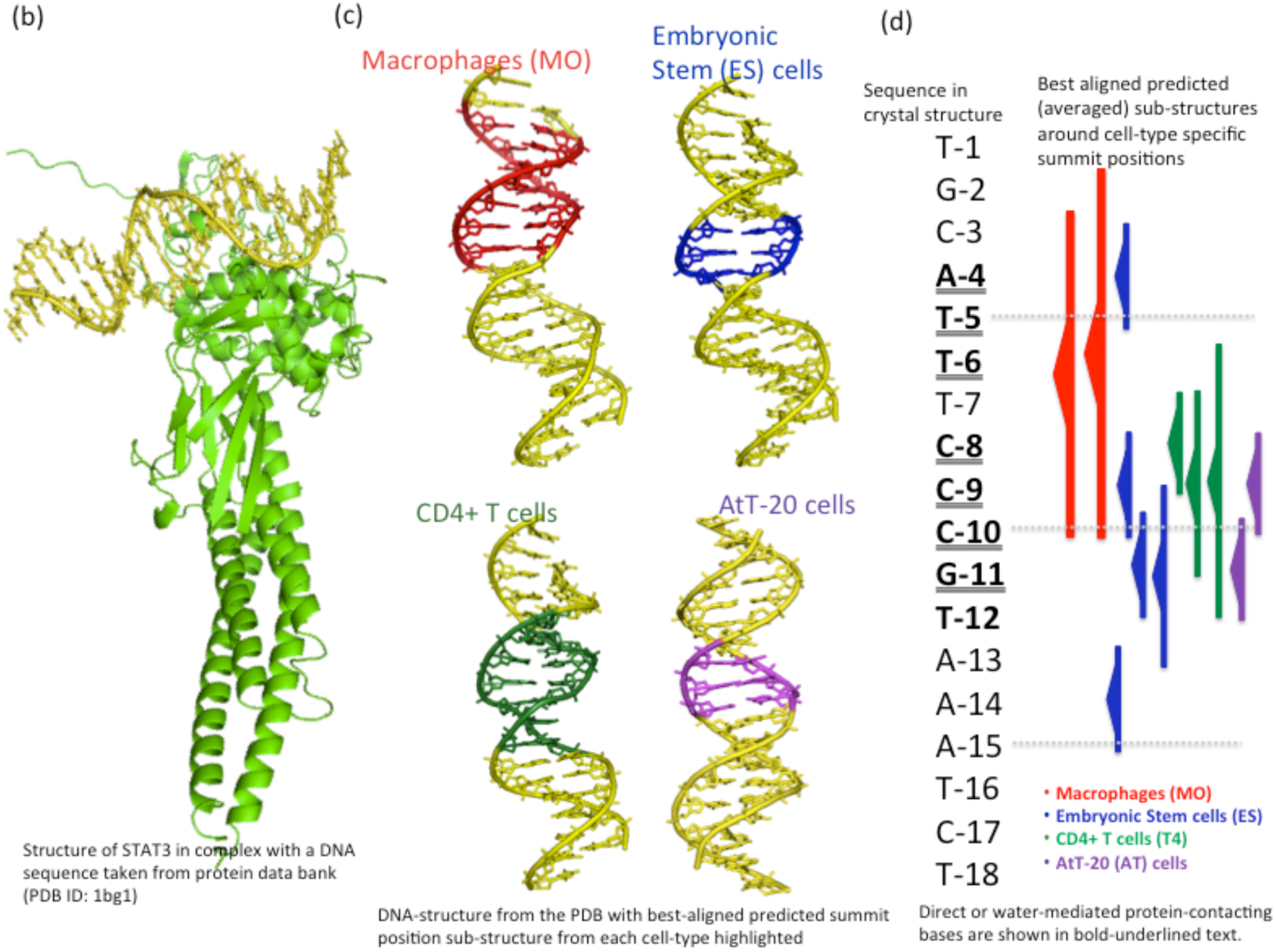
Structural regions containing summit positions from binding sites taken from different cell-types align to different positions of DNA in a crystal structure reported in the Protein Data Bank. Alignments are scored by the average of the 12 Pearson correlations between pairs of conformational parameters from predicted and observed structures. (a) Clustering cell-types and sequences in terms of alignment scores. Sequence names used are the ones observed in PDB and the number attached to the sequence at the end is the position of the first base in PDB at the start of an alignment. (b) Crystal structure of STAT3 complex with a DNA target sequence taken from PDB (presumed biological unit, PDB ID: 1bg1) (c) DNA molecule extracted from crystal structure and the best aligned predicted region of the genomic targets from four different cell types (d) Same information as in (c) mapped to the sequence observed in PDB.

## TF classification by target chromatin state

One of the mechanisms for context-specific activation of target genes by TFs is based on cooperative activation such that two or more TFs recognize their binding sites on promoter DNA sequence [29]. Cooperative recognition may follow either a prior formation of protein-protein complex or may proceed in a sequential manner whereby one TF opens the chromatin structure to enable the other to bind, the latter having no ability to binding closed chromatin state [15, 30]. Decoding these sequential patterns of cooperativity is critical to understand the molecular basis of gene expression in the genomic context. Recently, three categories of TFs namely *Pioneer Settler* and migrant have been defined based on the kind of target chromatin structure recognized by them. Sherwood et al. have provided a comprehensive characterization of TFs into these three groups using DNase-Seq profiles [15]. The *pioneer* TFs, are proposed to be distinct from others as they can occupy and are capable of opening the previously closed chromatin. After a *pioneer* opens the chromatin, TFs belonging to the second group called *settlers* bind to their targets[31] followed by a third group called *migrants*, whose member TFs do not bind independently but instead require co-factor interactions for DNA binding. Despite providing profound insights into TF recognition of various chromatin states, it is unclear what structural features of TFs or DNA determine its above characterization. We investigated whether the features derived from the intrinsic DNA conformational dynamics of TF targets (predicted by DynaSeq) can be useful to characterize them into these groups.

We started with the analysis of mouse ES cell DNase-Seq data described in Sherwood et al. Genome-wide TF-bound sequences were aligned by the first base of their respective motif start positions and computed the conformational ensemble signatures for each binding site followed by averaging of signatures for each TF (Figure 6). We investigated if there exists a common intrinsic conformational dynamics signature in the binding sites of these TF groups and if so, whether this signature is located on the putative binding-site, its flanking regions or both. As a first step, we asked whether the ensemble-based signatures of individual TFs within each of the three factor groups (*pioneer*, *migrant* and *settler*) correlate better within the group or across these groups. Figure 6(a) and 6(b) show the Pearson’s correlations between these ensemble signatures. Position-wise inter-group and intra-group TF-TF correlations within a 10-base window clearly show that the ensemble profiles at binding sites would not be able to distinguish between TF groups. However, ensemble profile similarities sharply increase away from the binding sites both for inter- and intra-group comparisons. This may not be surprising given that the distal regions of all TFs may represent similar chromatin locations, leading to similar conformational dynamics. What is remarkable, however, is that the intra-group correlations are consistently better than the inter-group correlations for the entire range of 200 base distances examined in this analysis. Figure 6(b) confirms that these differences are not caused by the presence of a few TFs with similar motifs, but present a more consistent trend. Heatmaps showing a comparison of predicted conformational dynamics, also demonstrate that the motif-depleted regions are powerful indicators of TF-TF similarities within a group Figure 6(a). Indeed, the two heatmaps in Figire 6(a) also show that *pioneer* and *settlers* are more similar to each other than each of them is to the *migrants*. This trend is in good agreement with the original definitions of *migrants*, which show weak activities of their own and requires other co-factors for binding. However, a remarkable outcome of this analysis is that the TF-binding classification by the chromatin state is encoded in the conformational ensembles of distal binding-site-flanking DNA sequences and not in the TFs or the sequence-specific TF binding sites themselves. The lack of a common ensemble signature between intra-group TFs may reflect sensitivity to individual aligned positions. Since each base position within the binding site contributes differently to binding in each TF, a common signature for several TFs cannot be defined. However, binding site-flanking regions are less position-specific and hence their TF-TF relationships are preserved. Irrespective of this possibility, the fact remains that distal binding site flanking regions carry a strong signature of the group to which a TF belongs.

**Figure 6.**
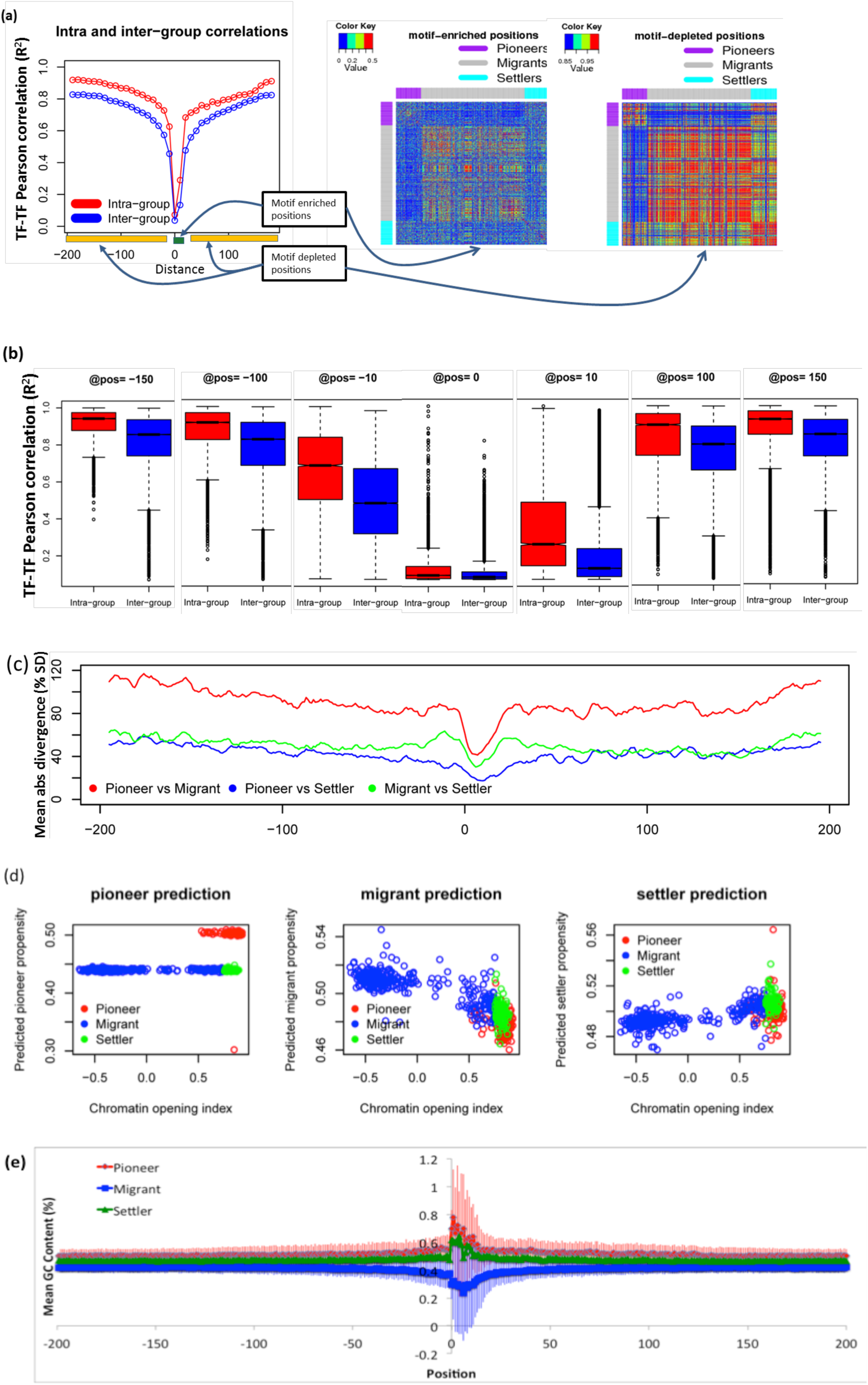
(a) TF-TF correlations between 10-base windows ensemble profiles. Motif-enriched positions show poor correlations between or within a TF group presumably because the importance of individual positions is highly TF-specific and TF-TF alignments distort any potential similarity. On the other hand motif-depleted regions show strong intra- and inter-group correlations, as distant regions become more and more similar to one another. However, intra-group TF-TF correlations are significantly better than inter-group correlations, especially in motif-depleted regions. Heatmaps show how TFs within the three groups of factors correlate with one another (b) Detailed distributions of the data for selected positions in (a), which confirms statistical significance of difference. (c) pairwise mean absolute divergence between factor groups, showing pioneer/migrant comparison is most significant as their mean ensemble populations differ from each other by about 100% of standard deviation. (d) Prediction performance of binary classifiers based on multiple linear regressions of ensemble-based features. Even though, the exact performance shown in this plot may be somewhat exaggerated due to redundancy between training and test data sets (similar TFs being present in the data), a relationship between conformational dynamics and pioneer behaviour detected in (a-c) is well supported (e) Average GC-content in the binding sites of each factor groups (standard deviations across TFs in pioneer and migrants shown as error bars). There is a significant GC-content enrichment in *pioneers* and *settlers*, but the same is accompanied by large inter-factor variation

Furthermore, ensemble signatures are informative of TF-group can also be seen from the pair-wise mean absolute divergence of ensemble populations at each alignment position (Figure 6(c)). As shown in Figure 6(c), pioneer and migrant factors are the two most distinct TF groups where the ensemble populations in each bin differ from each other by a modest 40% of standard deviation at the binding sites but by as much as 80-100% in their flanking regions. Other pairs of TF groups also show similar patterns of divergences albeit with a much lesser magnitude. Since on the average each single base position carries strong pairwise divergence, a cumulative signature comprising multiple positions is expected to be a powerful predictor of TF group. To quantify the ability of ensemble profiles of sequence regions (+/-200 bases of motif start position), we developed bootstrap cross-validation linear regression models with the aim to predict one TF group versus the rest in a binary manner. These models were developed for 5-base windows at each position and then averaged for the entire range to obtain a single value or “propensity” of a TF to belong to a particular group. Figure 6(d) plots the propensity scores against the chromatin-opening index (taken from Sherwood et al.[15]). (Complete results are in Supplementary Table ST5). Remarkably, we observe near perfect classification in predicting pioneer TFs, as all-but-one TFs were correctly identified as belonging to the pioneer group. Migrant TFs were also identified but with a much reduced accuracy (Figure 6(d)). Finally, settler TFs could not be predicted using this strategy, because their ensemble properties are in the ranges between migrants and pioneers, making a linear regression model unsuitable to classify them. More complex computational models are likely to show a better performance for settler prediction. Current data are sufficient to demonstrate that the conformational dynamics of the three groups of TFs significantly differ from one another and that these differences are most obvious at distal binding site-flanking regions instead of the binding sites.

In an attempt to find useful features at the binding sites, which could distinguish between the three groups of factors, we identified GC-content to be of interest. Both pioneer and settler factors show high GC contents in their binding regions which are depleted in the case of migrant factors (Figure 6(e)). However, as shown in this figure, despite differences in their mean values, TF-TF variations in GC content (revealed by standard deviations) are too high to consider them for predicting a TF group from this property. Additionally there is little difference between pioneers and migrants, whereas our classification model can accurately discriminate between these two groups. Conformational dynamics *ensemble features* therefore provide a much more comprehensive tool for TF group characterization.

## Discussion and Conclusions

The exact nature of conformational dynamics in TF recruitment, target search and complex stabilization is poorly understood and the role of binding site proximal and contiguous regions at genomic scales has not been reported before. Indeed, exactly how TFs scan the genome to find their binding sites is poorly understood [1]. Conformational dynamics of genomic targets of TFs has been elusive partly due to the lack of methods to perform large-scale simulations. With this work, we attempt to bridge this gap and show that predicted conformational dynamics provide important biological insights into TF recognition of its genomic targets.

TFs -exemplified by STAT3 in this work-show highly redundant sequence specific DNA binding [32] yet at the same time, they can exhibit highly specific cell-type activity. Moreover, the same family of TFs can be *pioneers*, *settlers* or *migrants* with no obvious link to their DNA-binding sequence. Here we show that the DNA regions much larger than the well known TF binding sequence motifs encode shape and specificity information for TFs, indicating that the genomic DNA is not just a ‘passive observer’ of TF binding. Instead, TF-DNA interaction is a mutual event between the DNA sequence and the TF, which act in unison to bring about a specific biological activity. This also reiterates the significance of allostery and cooperativity in protein-DNA recognition as implied from our previous works [29, 33-35] The allosteric effect in DNA targets in the recognition process is a subject of great interest [33-39]. Even though not reported for STAT3 targets before, our analysis suggests a role for allosteric control in STAT3 target recognition. The observation of four distinct regions around a binding site revealed by our analysis tempts us to think of them as composed of “*direct recognition*”, “*binding site stabilizing*”, “*recruitment and steering*” and “*background*” properties. However, additional data is needed to establish a functional association of these modular regions and to establish whether such modularity is universal to all TFs.

Binding site information away from the summits in STAT3 data are derived from ChIP-Seq and one could argue this to be an artifact caused by noisy mapping of summit positions. We believe that the number of binding sites is large enough to remove that noise. The clear separation between selected regions, existence of relatively sharp boundaries between them, and large changes in the slopes of piece-wise regression lines, suggest that distinct recognition regions do exist. The same has also been supported by a limited subset of data which contains high quality motifs, even though the sharpness witnessed in global data are somewhat blurred.

DNase-Seq data for *pioneer*, *settler* and *migrant* factors are more reliable in terms of the exact location of binding sites due to the experimental resolution and because it is pre-aligned by sequence motifs. Here also we established a role for conformational dynamics of binding site distal regions. Taken together, we believe that TF recognition in the genome is strongly influenced by conformational dynamics of much longer regions than previously assumed.

In conclusion, in this work, we present a novel approach to predict sequence dependent DNA-conformational ensembles directly from sequences, which does not require detailed simulations of their structures. Models were trained and cross-validated on MD simulations of all unique tetra-nucleotides and perform well in various evaluation tests. Two complementary model systems representing genome-wide binding preferences of TFs in a cell type-specific and mode of TF-binding were analyzed. In the first, genome-wide binding preferences of STAT3 across different cell-types could be modeled using features derived from predicted conformational ensembles with accuracies better than sequence-only information. Superiority of DNA conformational dynamics over its static structures in accurately modeling STAT3 binding data is also established. In the second, TF characterization as *pioneer, settler* and *migrant* transcription factors is explained with high accuracy by conformational dynamics of regions distal to, but not at the binding sites. Together, these results suggest the cooperation of much larger chromatin regions in transcription factor binding activities than realized so far.

## Methods

### Definitions

*Base-step* at a position *i* in a DNA sequence is defined as the pair of nucleic acid bases at position *i* and its 3’ neighbor *i+1*.

*Conformational parameters* here correspond to 12 unique helical and base-step parameters: *shear, stretch, stagger, buckle, prop-tw, opening, shift, slide, rise, tilt, roll* and *twist*, as provided by the *analyze* option of *3DNA[16]*. These are summarized in Supplementary Table ST1. While, helical parameters are straightforward to assign to each base position, the base-step parameters, strictly defined for a pair of bases rather than a single base have been also assigned to each base position *i* in this work.

*DNA conformational ensemble* or just *ensemble* refers to the observed or predicted population density distribution of a single base (and by extension for a DNA sequence) in pre-defined bins (ensemble bins) representing ranges of base-step conformational parameters. One ensemble contains population density data from 12 conformational parameters and is computed from snapshots of the 3D structure in a Molecular Dynamics (MD) trajectory or predicted by a trained model directly from the sequence. There are five bins for each conformational parameter, whose ranges are defined by equal frequency distribution in the global data (see Supplementary Methods SM2).

*Window size:* This term has been used to indicate the sequence-window being under consideration in a given model and its actual value depends on the context. For example, the final sequence-based SVM model *DynaSeq* is based on a 5-base window. Once the ensemble is predicted by *DynaSeq*, predicted features with a sequence window of a larger size are often concatenated, and the same is specified in the corresponding results accordingly.

### Principle components

Sequence data at each base position is encoded by a four dimensional binary vector, in which a non-zero value represents the identity of the base located at that position. Conformational ensembles is encoded by 60 dimensions (dynamics) and its averaged structure is encoded by 12 dimensions, as described above. Principle components of the dynamics and structure are computed by first predicting a them in their native dimensionality (60 and 12 respectively) for all the unique 6-mers (for consistency with DNA Shape, needed in some comparisons) and then computing their principle components using the R-package *prcomp (*www.r-project.org*)*.

### STAT3 targets data sets

*Mapping and alignments:*

Genome-wide STAT3 binding data from ChIP-Seq experiments in four cellular and contexts, were used for these comparisons and obtained from our recent works, where the complete procedures for mapping and peak calling are described in detail[14]. Sequences were aligned by their motif centers detected by MEME-CHIP program [40]. In each case three motifs were detected and sequences containing these motifs were compiled for motif-aligned analysis.

*Cell type names* are abbreviated as MO for macrophages, ES for embryonic stem cells, T4 for CD4+ T cells and AT for At-T20 cells.

ChIP-Seq *summit* or simply a summit position in TF binding site data is defined as a genomic position, where a local maxima of the number of aligned sequence reads is observed. This position is believed to be the midpoint of a putative binding site and multiple *summits* each corresponding to an independent putative binding site are observed in a single ChIP-Seq experiment.

*Control genomic sequences*: Genomic background sequences for each set of STAT3 binding sites are equally sized sequences centered at the random control tags taken from the corresponding ChIP-Seq library data of the four cell types considered in this work.

### Pioneer, migrant and settler factor data sets

List of pioneer and migrant factors was taken from the supplementary tables provided in[15]. Binding site coordinates data were taken from the related online resource located at (http://piq.csail.mit.edu/data/v1.3.calls/140906.mES.calls.tar.gz) from the same authors (data was downloaded on October 1, 2014 and has been reorganized on the authors’ website since then). Final list of pioneer and migrant TFs used in the current work is shown in Supplementary Table ST5, along with chromatin opening index from the original source and pioneer propensity score predicted by our method.

### Feature enrichment analysis

Predicted population in a bin or the averaged value of a conformational parameter represents a feature, which may or may not contain information about sequence specificity. Predicted conformational ensemble for each position in the data set is expected to have a score of 0.20 (there are five equal probability bins) and this reference can in principle be used to determine specificity/enrichment of ensemble features in ChIP-Seq data sets. However, actual genomic sequences may have different nucleotide composition than those used in defining equal frequency bins and therefore signatures of target sequences are determined by comparing predicted ensemble populations between ChIP-Seq data with those for the random genomic ones. A specific divergence *D* for a between control and real ChIP Seq ensemble for a feature *F* is computed as:

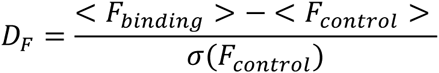

Here feature *F* refers to the population of a single conformational bin (comparison of dynamics), or averaged predicted value of conformational parameter (comparing of structures), <x> and σ(x) respectively denote the average and standard deviation of x. Thus 60 and 12 divergence scores at each position (distance from the summit) respectively characterize how the dynamics and structure of observed targets differs from control.

Assuming the random sequences to be non-binding, we trained multiple linear regression (MLR) classifiers to distinguish between binding and non-binding sequences. Inputs to these regression models are predicted ensemble populations for all nucleotide positions within a fixed window size and expected output is a score representing the probability of the sequence being binding or not. In all cases training is performed over 80% of the sequences, selected randomly and tested on the remaining 20% data. Five such cycles of sampling are carried out and performance scores are averaged. In order to make a fair comparison of dynamics-based models with sequence-only and structure-only models, similar independent MLR models were generated for each of these feature sets as inputs. MLR coefficients were obtained using GLM package in R programming environment and prediction scores were for the test data were computed using in-house dedicated scripts, all implemented in R[41].

### Prediction of pioneer, migrant and settler propensity

Prediction of pioneer, migrant and settler propensities for a TF is performed in two steps i.e. bootstrapped regression models trained over features from small sequence sliding windows (5-base window in all cases) followed by averaging over all window positions over all bootstrapped cycles. For example, when predicting pioneer propensity of a TF, first a class label of +1 or 0 is assigned to all pioneers (positive class) or non-pioneers (control class) respectively. A 60 dimensional ensemble signature is computed for each TF at each position (motif start +/-200) by taking the average ensemble from all its binding sites. Using a 5-base window at a specific alignment position for all TFs a regression model is trained in a bootstrapping cross-validation i.e. training 70% randomly sampled data and finding predicted score for the remaining 30%. Sampling and training are repeated for 100 bootstrap cycles and the sliding window is moved for all alignment positions (200*2+1 minus 4 terminal positions). All predictions from all bootstrap and sliding window predictions are averaged to obtain a single propensity score. Three such independent propensity scores corresponding to pioneer, migrant and settler as positive class respectively for each TF are obtained in this way.

## Authors’ contributions

SA conceived and designed the main components of this study and implemented it together with MA. APH prepared the mapped sequence reads of STAT3 targets and controls, and helped in the analysis of STAT3 targets together with DMS. HK provided MD data and helped in analyzing other results. SA, MA and APH prepared the manuscript with discussions and critical comments from RN, DMS, HK and KM. All authors participated in discussions on all aspects of the manuscript and read, improved and approved the final version of the manuscript.

## Acknowledgements

This work has been supported by a grants-in-aid (kaken-hi #15K00419) program of Japan Society for Promotion of Science of Japan to SA.

This work has also been supported by Grants-in-Aid for Scientific Research from the Ministry of Education, Culture, Sports, Science and Technology (MEXT) of Japan [25116003 to H.K] and Platform for Drug Discovery, Informatics, and Structural Life Science from the MEXT to HK.

This study was in part supported by Grants-in-Aid for Scientific Research from the Ministry of Education, Culture, Sports, Science, and Technology (Grant Numbers 25430186 and 25293079) and from the Ministry of Health, Labor, and Welfare to K.M.

This project has been funded in whole or in part with Federal funds from the National Cancer Institute, National Institutes of Health, under contract number HHSN261200800001E. The content of this publication does not necessarily reflect the views or policies of the Department of Health and Human Services, nor does mention of trade names, commercial products, or organizations imply endorsement by the U.S. Government. This research was supported (in part) by the Intramural Research Program of the NIH, National Cancer Institute, Center for Cancer Research.

